# Molecular Dynamics Trajectory Analysis of Permeation (MDTAP): A tool to analyze permeation events across membrane proteins

**DOI:** 10.1101/2024.04.12.589220

**Authors:** Palur Venkata Raghuvamsi, Sruthi Sundaresan, Thenmalarchelvi Rathinavelan

## Abstract

**Background and Objective:** Molecular dynamics (MD) simulations are indispensable and versatile in capturing the time-dependent conformational changes of biomolecules to shed light on the concomitant biological processes. MD is used to provide critical mechanistic insights into the transportation of solvent/solute/drug molecules across protein channels embedded in a membrane bilayer. The huge size and volume of the MD trajectories of a membrane-embedded system provide challenges in the analyses of membrane permeation events. Thus, a software, Molecular Dynamics Trajectory Analysis of Permeation (MDTAP), is presented here to analyze the permeation events across membrane-embedded proteins and nucleic acids automatically.

**Methods:** A software is developed here to automatically detect the permeation events across the channels irrespective of their shape and size and the type of solute molecules from the MD trajectories. MDTAP employs bash scripts to fetch information about the permeation, residence time, and diffusion of the molecules of interest in a Linux/Mac-based environment. The source code of MDTAP is freely available to the public, along with installation and usage information on GitHub (*attached as supplementary for the review process and will be made accessible to the public through the following link upon acceptance for publication:* https://github.com/MBL-lab/MDTAP).

**Results:** The efficiency of MDTAP is demonstrated here by considering the MD trajectories of 2 water-conducting channels as test cases: *E. coli* outer membrane protein Wzi and *E. coli* Aquaporin Z. The dimensions of the channels and their capacity to accommodate and conduct water, the number of permeating water molecules along with the path traced and time taken to cross the channel is validated.

**Conclusion:** In summary, the graphical representation of the time-dependent behavior of the solute/solvent permeation events corresponding to an MD trajectory in MDTAP allows the user to easily visualize the mechanism of permeation, including the localization of the permeating molecule (if any) and permeating path. Thus, MDTAP immensely reduces the difficult task of manually analyzing solute/solvent permeations from the bulk MD trajectories. Such a simplistic representation of permeation events across the protein transporters helps in the design of drug molecules to treat the associated diseases. Further, MDTAP is also designed to characterize the permeation events across artificial nucleic acid channels, considering their importance in recent times.

## Introduction

Molecular dynamics (MD) simulation has emerged as a computational microscope to capture the time-dependent conformational dynamics of biomolecules [1, 2]. Since its flourishment in the 1970s, the efficacy of MD simulation techniques in providing structural insights about biological phenomena has been well demonstrated [3-9]. One such example is the translocation of small molecules like water, ions, or other solute molecules, which play a pivotal role in maintaining cellular homeostasis across membrane-embedded protein channels [10]. For instance, the mechanism of water conduction by aquaporins, a water-conducting membrane protein that regulates the amount of water in cells across all the kingdoms of life [11], was elucidated in greater detail by MD simulations [12, 13]. Similarly, MD helped understand the gating mechanisms of mechanosensitive [14], VDAC [15], and potassium channels [16], *etc*. [17, 18]. MD also was helpful in understanding the solute conduction mechanism across artificial nucleic acid channels [19].

There are many tools to map the transmembrane channels, like CHAP [20], HOLE [21], CAVER [22], and MOLEonline [23]. Interestingly, the PerMM tool differs from the others [20-23] as it provides the permeation coefficient and the translocation pathway of drug-like molecules through artificial and natural membranes [24]. However, to the best of our knowledge, there is no tool available to capture the solute/solvent conduction events across a membrane protein from the vast volume of MD trajectory in an automatized fashion irrespective of the nature of the pore shape, pore size, and directionality of transportation (*viz*., unidirectional (extracellular to intracellular or vice-a-versa) or bidirectional). To this end, an MD analysis tool, **<u>M</u>olecular <u>D</u>ynamics <u>T</u>rajectory <u>A</u>nalysis of <u>P</u>ermeation** (MDTAP), is developed here to capture and quantify the permeation events across protein and nucleic acid channels. MDTAP offers flexibility for the user to define the molecule of interest whose permeation across a protein or nucleic acid channel can be tracked using the PDB structures generated from the MD trajectory irrespective of the mode of conduction (*viz*., a single molecule in narrow channels or bulk molecules in wider channels which may either be unidirectional or bidirectional) of water/ions/solute/small molecules.

MDTAP uses a distinct approach to facilitate the capture of molecules transported across a channel having a straight or tortuous diffusion path. MDTAP also effectively captures solvent/solute/drug molecule diffusion patterns adopted by the channel of the user’s interest and calculates the flux and coefficient of permeation and conductance (in the case of ion channels). The source code of MDTAP is freely available without any registration to the public, along with the installation and tutorial on GitHub (*attached as supplementary for the review process and will be made accessible to the public through the following link upon acceptance for publication:* https://github.com/MBL-lab/MDTAP).

### Statement of significance

- Conformational dynamics of biomolecules are essential for the understanding of mechanisms underlying human diseases
- Molecular dynamics (MD) simulation acts as a computational microscope
- MD enables the capturing of molecular conduction events across membrane channels
- A methodology is developed here to ease the characterization of permeation, automated and implemented as a Linux/Mac-based software, MDTAP
- Validation is done using channels having tortuous and straight diffusion paths

## Methods

### Development of a methodology to calculate permeation events

Water/ion/solute/small molecules are considered permeating only if a complete translocation occurs through a channel from one side to the other. The permeation path of the molecule(s) may not always be a single file. One such example is given in Figure 1A, which is a water-conducting [25, 26] outer membrane channel of *E. coli*, Wzi [27]. Besides, this protein channel has multiple entries and exits located at both periplasmic and extracellular sides. Due to multiple entry and exit points, a water molecule entering the Wzi channel does not necessarily have to cross the channel during the MD simulation time. Instead, it may simply diffuse in and out without crossing over to the other side of the channel or even stay within the Wzi channel during the simulation time due to its wider and tortuous nature. Thus, it complicates the understanding of water permeation mechanism(s). In order to capture the molecular permeation events across a channel of any type (*viz*., straight or tortuous and single or multiple entries/exits) in an automated fashion from the MD simulation trajectories, a distinct approach is devised here.

**Figure 1.**
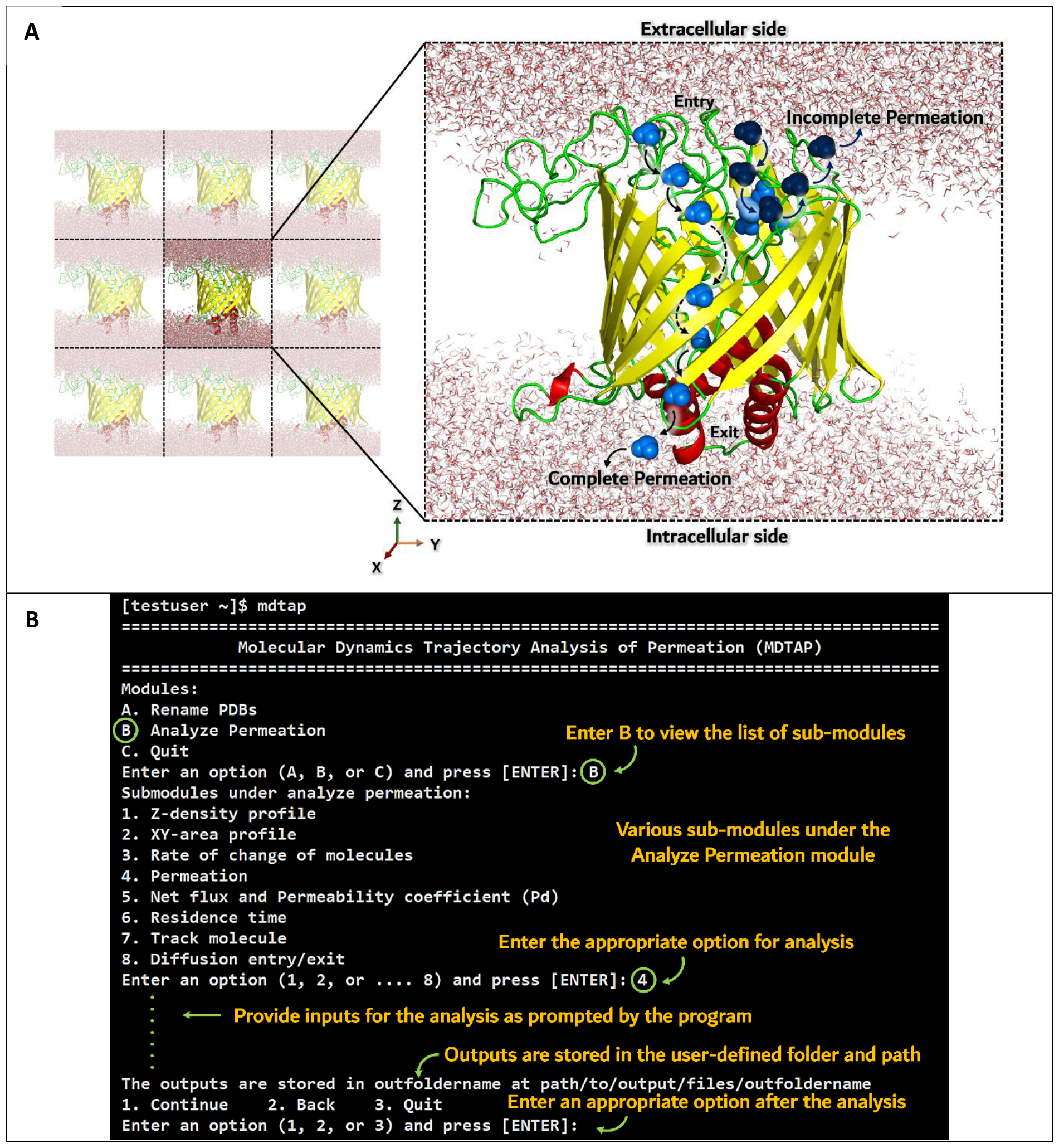
Methodological development of MDTAP. A) A schematic describing true/complete and non/incomplete permeation events in an automated fashion is illustrated by considering the water-conducting *E. coli* Wzi protein (cartoon representation, PDB ID: 2YNK) as an example. Wzi in a periodic boundary condition is shown on the left. For the sake of clarity, POPE and POPG (3:1) membrane molecules are not shown. The water reservoir on either side of the channel is shown in line representation. While the light blue colored water molecule shows the complete permeation event, the dark blue colored water molecule shows an incomplete permeation. B) Snapshot of MDTAP software interface.

The methodology is devised to initially fetch the solute/solvent/ligand molecules present within the channel (aligned along the Z-axis) dimension in the given MD trajectory (**Figure 1A**). Subsequently, the frequently occurring molecules are shortlisted based on the number of occurrences of the molecule(s) of interest inside the channel. Next, based on the vicinity of the shortlisted molecule(s) to the two different sides of the channel Z-axis (for example, the top or bottom of the channel) with respect to time (*viz*., entry or exit time), the direction of permeation event is decided. Finally, ensuring that the difference between the lowest and the highest Z-coordinates of the molecule is greater than or equal to the channel dimension during the permeation event confirms that the molecule(s) of interest crosses the channel. This method also facilitates straightforwardly identifying the bidirectional permeation events irrespective of the locations of the molecule of interest (*viz*., top or bottom of the channel). Further, the molecules/ions that don’t completely cross the channel would automatically identified as non-permeating. This approach also eliminates the chance of reporting false permeation of the molecule of interest. This scenario occurs in the pre-filled channels, wherein the molecule (here, water) is already present inside the channel (happens during the equilibration part of the simulation). Thus, a molecule that is already located inside the channel will not be considered a permeating one.

### Implementation of the methodology as a permeation analysis software

The methodology is implemented as a Linux/Mac-based software named “MDTAP: Molecular Dynamics Trajectory Analysis of Permeation” and subsequently validated. The software executes this methodology in a bourne again shell (bash) environment. The software has multiple interactive modules, as detailed in the further sections, along with the input and output information. Since some modules generate output graphs for the user’s preliminary inspection, the system executing the software should have gnuplot installed and can be accessed by the name “gnuplot”.

### Rename PDB and exit modules

The interface of MDTAP software is given in **Figure 1B** and **Table S1**, wherein the user has to first execute the “Rename PDBs” module (invoked by ‘A’ or ‘a’) to rename the PDBs (generated from the trajectories of any MD simulation package) to make them accessible to MDTAP (**Table S2**). This module renames the PDBs in sequential order with the prefix “step_” and “.pdb” extension (*viz*., step_1.pdb, step_2.pdb, step_3.pdb, *etc*.). Note that the user can leave the software interface by using options ‘C’ or ‘c’ of the main module.

### Analyze Permeation Module (APM)

Since parameters like permeation coefficient, pore dimension, diffusion events, *etc*., are essential to characterize the permeation of a solvent/solute/small molecule(s) across a channel, the second module of the software “Analyze permeation” (henceforth, APM; invoked by typing ‘B’ or ‘b’) has eight submodules as described in **Table 1** and **Figure S1**. While all the modules would be very helpful in understanding the molecular transport across the reservoirs on either side of a channel, modules 6-8 would help analyze a single molecule’s transport across a channel [28]. Notably, all the APM submodules assume that the channel is aligned with respect to the Z-axis of the Cartesian coordinate system. **Figure 2** summarizes the outputs generated by the submodules of the “Analyze permeation” module by considering the previously published *E. coli* Wzi trajectory [25] as a test case. To use modules 1-4 and 6-8, the user must first specify the location of the renamed PDBs. Similarly, modules 1-4 and 8 require the residue name and one of the atom names of the molecule of interest (for example, “OH2 TIP” represents water oxygens OH2, having a residue name TIP). Modules 1-4 and 6-8 allow the user to define the starting and ending PDBs along with the number of PDBs to be skipped for the analyses (*viz*., for start, end, and skip values of 11, 21, and 2, respectively, the PDB files considered for the analyses would be step_11.pdb, step_13.pdb, step_15.pdb, …step_21.pdb). These modules further require the time difference (in picoseconds) between the consecutive PDBs (**Figure S1**).

**Table 1.** List of MDTAP modules and their functionalities. The module number to be used in the software interface to invoke the appropriate module is given in column 1. Note that these modules do not need to be executed in a sequential order.

| APM Number | Module Name | Function |
| --- | --- | --- |
| 1 | Z-density profile | Identifies the variation in the population of permeating molecules along the channel axis (*viz.*, Z-axis) |
| 2 | XY-area profile | Identifies the aerial and spatial distribution of the molecule of interest projected to the XY-plane |
| 3 | Rate of change of molecules | Calculates the number of molecules present inside the channel with respect to time |
| 4 | Permeation | Identifies and lists molecules that undergo complete and incomplete permeation through the protein |
| 5 | Net flux and permeability coefficient ($P_d$) | Provides the net flux and the diffusion permeability coefficient ($P_d$) of the molecule of interest across the channel |
| 6 | Residence time | Calculates the time (in ps) that the permeating molecule resides in the conduction path |
| 7 | Track molecule | Tracks the path followed by the permeating molecule along the pore axis (Z-axis) as it passes through the channel |
| 8 | Diffusion entry/exit | Calculates the number of molecules that enter or exit through a diffusion plane defined by three amino acids |

**Figure 2.**
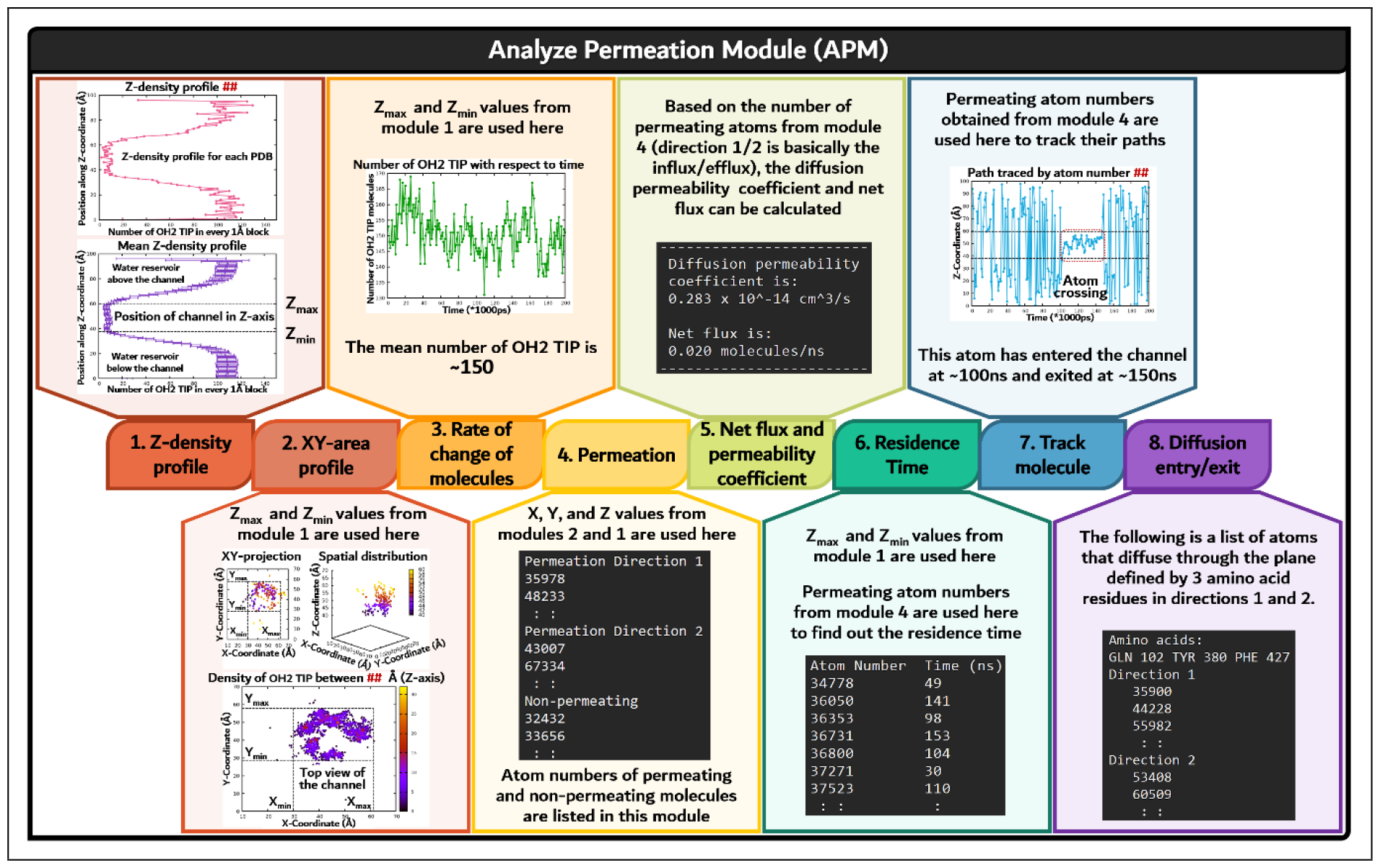
Schematic describing the outputs generated by the eight different submodules of the “Analyze permeation” module (APM) (See **Table 1** for details) by considering *E. coli* Wzi trajectory (200 ns) [25] as a case in point is shown. The channel’s position with respect to the Z-axis is obtained from the “Z-density profile” module and is used as the Z_max_ and Z_min_ values for the subsequent modules. Similarly, the “XY-area profile” module gives a clue about the channel’s position with respect to the XY plane. The list of atom numbers that are found to permeate across the channel is obtained from the “Permeation” module (*viz*., the atom numbers of those molecules that undergo complete or true permeation) and is used in the subsequent modules. The red hashtags (**##**) in the graph titles indicate that it is a representative output of the respective module.

Modules 1-4 and 6 require the dimension of the channel of interest (*viz*., channel limits). Thus, MDTAP automatically recommends the channel dimension (*viz*., X_max_, X_min_, Y_max_, Y_min_, Z_max_, and Z_min_) based on the minimum and maximum values of X-, Y- and Z-coordinates of the protein/nucleic acid channel. This automatically captures the permeation event(s) even if the channel extends beyond the inner-/outer-membranes (extracellular, periplasmic, or cytoplasmic sides). One such example is Wza, an *E. coli* protein that spans beyond the outer membrane bilayer [29] (**Figure S2**). The direct usage of channel coordinates for the permeation calculations further allows the user to use post-processed MD trajectories even in the absence of a membrane bilayer.

To facilitate the multichannel analysis, the software also allows the user to enter the segment and/or chain IDs of the channel of interest in modules 1-4 and 8. For instance, if there are four channels as in aquaporin, which are given seg ID names as ‘segA’, ‘segB’, ‘segC’, and ‘segD’ in the input PDB files, the user can specify ‘segA’ to analyze the permeation across the first channel. In the absence of a seg ID in the PDB files (like in a single channel), the user may skip this option by pressing the enter key. Similarly, the user can use the chain ID as an input, the absence of which in the PDB, the user may skip this option by pressing the enter key.

Unless otherwise mentioned, all other modules require similar input.

#### Z-density profile

The first module under APM is “Z-density profile”. This module provides information about the molecule’s conduction path and generates a graph showing the “Number of molecules in every 1 Å block vs. Position along Z-coordinate (Å)” as shown in **Figure S3A**. After fetching the appropriate information from the user (**Table S3**), the location of the transporting molecules along the Z-axis (*viz*., the channel axis) is generated for each 1 Å block by counting the number of permeating molecules. Note that, unlike the other modules, this module uses the maximum and minimum values of Z-coordinates of the entire system to fetch the Z-axis dimension of the channel. However, the user can also manually define the dimensions in which they are interested. This module at the end generates “Number of molecules in every 1 Å block vs. Position along Z-coordinate (Å)” plots and text files individually for each input PDB, as well as a time-averaged plot (along with the standard deviations) and the corresponding text file.

#### XY-area profile

The “XY-area profile” module provides insight into the localization of the permeating molecules in the XY-cross-sectional area of the channel. The input for this module is similar to the previous module (**Table S4**). However, to specify the channel dimension in the input, the user can get an idea about the Z-axis channel dimension from the plot obtained from the previous module and use it to define the upper and lower Z-axis cut-offs. This module generates a series of 2D plots to give the user a clue about the localization of the transporting molecule in the XY plane with respect to time (**Table S4**). Along with the 2D plots, the module also generates a series of 3D plots with respect to time by incorporating the Z-axis coordinate of the permeating molecule in the third dimension (for each input PDB). The module further generates a series of density plots (using all the input PDBs), which is the projection of the transporting molecule onto the respective XY plane for each 1 Å block defined along the Z-axis to precisely obtain the pore size (in the XY plane) of the channel (**Figure S3B**). An animated gif file is also created using all the density plots (**Movie 1**).

#### Rate of change of molecules

The “Rate of change of molecules” module estimates the number of molecules inside the channel with respect to the simulation time to get an idea about the capacity of the channel to accommodate the molecule of interest. Defining the X-, Y-, and Z-axis dimensions based on the plots generated by the previous modules is encouraged (**Table S5**). Upon fetching the necessary information, the module generates a text file and the corresponding “Time (ps) vs. Number of molecules” graph, which provides information about the molecules inside the channel (**Figure S3C**). Further, the module provides a text file with the volume occupied by the molecule of interest within the defined X, Y, and Z dimensions with respect to time in picoseconds (ps) by multiplying the number of molecules with the molar volume of the molecule of interest.

#### Permeation

The “Permeation” module helps to calculate the number of molecules permeating across the channel irrespective of directionality (**Table S6**). A “permeation event” is a complete translocation or transport of water, ions, or small molecules through the channel of interest from one reservoir to another. The outcome of this module is provided in three text files (Permeation-dir1.dat, Permeation-dir2.dat, and Non-permeating.dat) that list the molecules that are being conducted in two different directions and the molecules that are non-permeating. Note that the number of molecules crossing in one specific direction alone will be listed for a unidirectional channel. To further visualize the path traced by the permeating molecule, the user can also check the PDB files generated by this module for each atom number. A single PDB is generated for each atom number, which consists of the positions of the permeating molecule during its residence inside the channel (**Figure S4)**.

#### Net flux and Permeability coefficient (P_d_) calculation

Insights obtained from the “Permeation” module can be used to calculate net flux and permeability coefficients using the “Net flux and permeability coefficient” module (**Table S7**). The net flux is calculated by subtracting the total number of permeating molecules in one direction from the opposite direction (for bidirectional channels) during the simulation time frame. Such a net flux calculation from the simulation trajectories can be utilized to observe the channel conduction mechanism and determine the attainment of equilibrium, wherein the number of molecules translocating both sides should be equal. This is done by counting the number of water molecules crossing the channel in 2 different directions by considering the total number of permeating molecules listed as direction 1 and direction 2 in the “Permeation” module. The resultant number will be the net flux. For the unidirectional channels, the number of permeating molecules will be zero in one of the directions. The module also calculates the diffusion permeability coefficient (P_d_) as described elsewhere [30]:

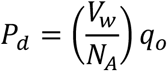

where, V_w_ is the molar volume of the molecule of interest, which is 18.07cm^3^ for water, N_A_ is Avogadro’s number, and q_o_ is the number of permeation events in one direction per unit of time. Due to the equilibrium conditions, the number of permeation events in both directions should be nearly similar for bidirectional conduction. Thus, the total number of permeation events is 2q_o_. In MD simulations, an equal number of bidirectional permeation events may not occur during the simulation time. Thus, an average of the permeating molecules is considered to calculate q_o_.

#### Residence time

The “Residence time” module calculates the residence time (in ps) of one or more molecules of interest inside the channel. This module requires information about the location of the input PDB files, the dimension of the channel, etc., along with a text file with a single column having a list of atom number(s) of the permeating molecule(s) (**Table S8**). The user can generate this text file after getting an insight from the “Permeation” module outputs. The generated two-column output text file provides the residence time corresponding to atom numbers listed in the input file (*viz*., atom number and residence time). The calculated residence time gives the time the permeating molecule(s) takes to cross the channel in either direction.

#### Track molecule

This module helps users to fetch the molecule’s permeation path in the Z-direction. This requires the same inputs as the “Residence time” module (**Table S9**) and generates a “Time (ps) vs. Z-coordinate (Å)” plot (**Figure S3D**) and text files for each permeating molecule. To trace the path taken by the permeating molecule in the Z-axis, the user can use the atom numbers generated by the “Permeation” module (**Figure S5**).

#### Diffusion entry/exit

This module provides an insight into the number of entries/exits of the molecule(s) of interest through a diffusion point. For this, a diffusion plane is defined, through which the molecule(s) of interest must pass to be considered as a diffusion [25] (**Figure S6**). A diffusion entry or exit plane is defined by calculating the center of mass (cm) (*viz*., X_cm_, Y_cm,_ and Z_cm_ indicated by O in **Figure S6**) for three user-defined amino acids located at the entry/exit point (**Table S10**). A local XY-plane (diffusion plane) is subsequently defined with upper and lower bounds of (X_cm_ + 5 Å, Y_cm_ + 5 Å) and (X_cm_ − 5 Å, Y_cm_ − 5 Å), respectively. The entry of the molecule of interest into the channel, which is aligned along the Z-axis, is considered as a diffusion only when the molecule has a Z-coordinate greater than Z_cm_ (let us say, Z_2_ as indicated in **Figure S6**) at the n^th^ ps and subsequently falls within the defined XY-plane having the Z-coordinate value between Z_cm_ and (Z_cm_ − 3.5 Å) (Z_1_), and also crosses Z_1_. Similarly, the diffusion exit is calculated in such a way that a molecule that has Z-coordinates lesser than Z_cm_ (let us say, Z_1_) at n^th^ ps and subsequently falls within the defined XY-plane having the Z-coordinates value between Z_cm_ and (Z_cm_ + 3.5 Å) (Z_2_) and further crosses Z_2_.

## Results

The implementation of the MDTAP methodology is validated by considering two water-conducting channels as the test cases. The first test case is an *E. coli* outer membrane protein Wzi, which conducts water in a passive bidirectional manner and has a tortuous path. The second test case is *E*.*coli* aquaporin Z (Aqp-Z). In contrast to Wzi, Aqp-Z is a single file mode unidirectional water transporter with a narrow pore.

Wzi represents the channels or transporters with a wide pore, wherein the conduction takes place in bulk, and the molecule(s) freely diffuse across the protein channel. An earlier MD study has shown that Wzi conducts water from the periplasmic side to the extracellular side and vice-versa [25]. These calculations were done by manual inspection of the trajectories, which is a laborious task. Here, it is shown from the last 200 ns trajectory of the published Wzi simulation [25] that MDTAP can automatically fetch the list of permeating molecules (**Figure S4**) and the path traced by them in the Z-direction (**Figure S5**). Thus, the last 200 ns trajectory of the published Wzi simulation is taken to test MDTAP’s automation in calculating the P_d_ and permeation events and is validated. Similarly, the 50 ns trajectory of Aqp-Z is also used to calculate the permeating events, and the results generated are shown in **Figure S7**. In summary, the results are in conformity with the published values, thus validating the MDTAP implementation.

### Limitations

The current version of MDTAP supports the trajectories in PDB format as it is universal.

## Conclusions

Molecular dynamics simulation plays a crucial role in understanding the dynamics of various membrane-embedded natural and artificial channels. However, manually analyzing the conduction mechanism from the MD trajectory is very time-consuming. Thus, a user-friendly software package called MDTAP is developed here to analyze the MD trajectories automatically (in PDB format). The program modules contained in MDTAP are flexible enough to take the user-defined inputs specific to the system under study. To the best of our knowledge, MDTAP is a one-of-a-kind analysis tool dedicated to characterizing permeation events across channels as it uses a methodology based on the channel dimensions to monitor the permeation of the molecule of interest. Irrespective of the mode of conduction of the molecule(s), *viz*., single file or bulk mode of translocation (**Figure S8**), MDTAP successfully captures the conduction of molecules across the channel as demonstrated by considering *E. coli* Wzi and *E. coli* aquaporin Z as examples. Thus, MDTAP is a dedicated package that could analyze the MD data specific to transporters.

## Code availability

The codes are deposited in GitHub and can be accessed through https://github.com/MBL-lab/MDTAP (*attached as supplementary for the review process and will be made accessible to the public through GitHub upon acceptance for publication*).

## Acknowledgments

The Computer Center, IIT Hyderabad, is acknowledged for its support. SS and RV thank MoE for fellowships.

## Ethical Approval

Not Applicable

## Author contribution

RV developed the methodology and implemented the software. SS fine-tuned the implementations and validated the results. RV, SS, and TR wrote the manuscript. TR designed and supervised the project.

## Supplementary figures and movie

**Figure S1.**
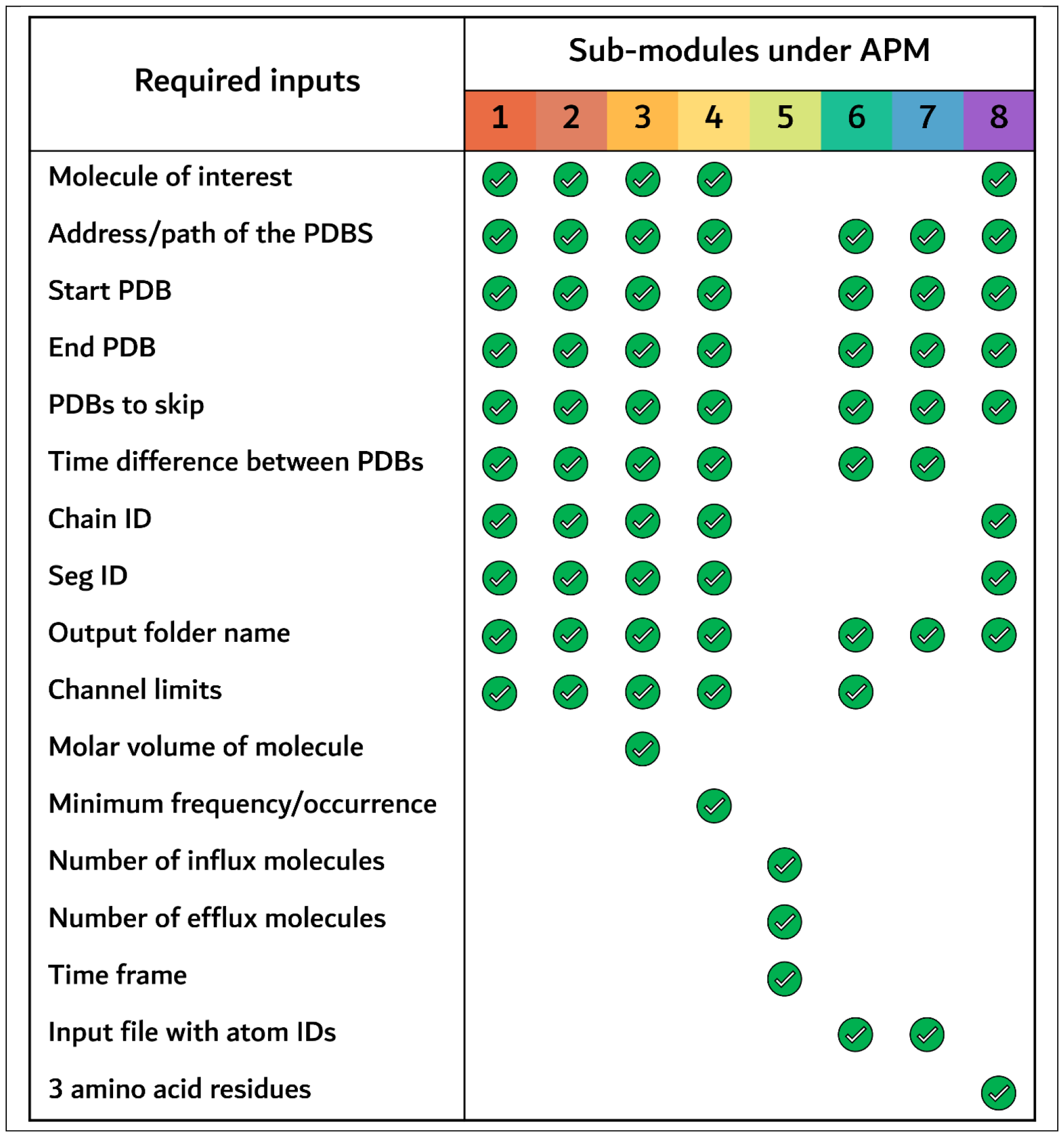
Inputs required by each submodule listed under APM. The submodules are indicated by their respective APM numbers, as in **Table 1** of the main text.

**Figure S2.**
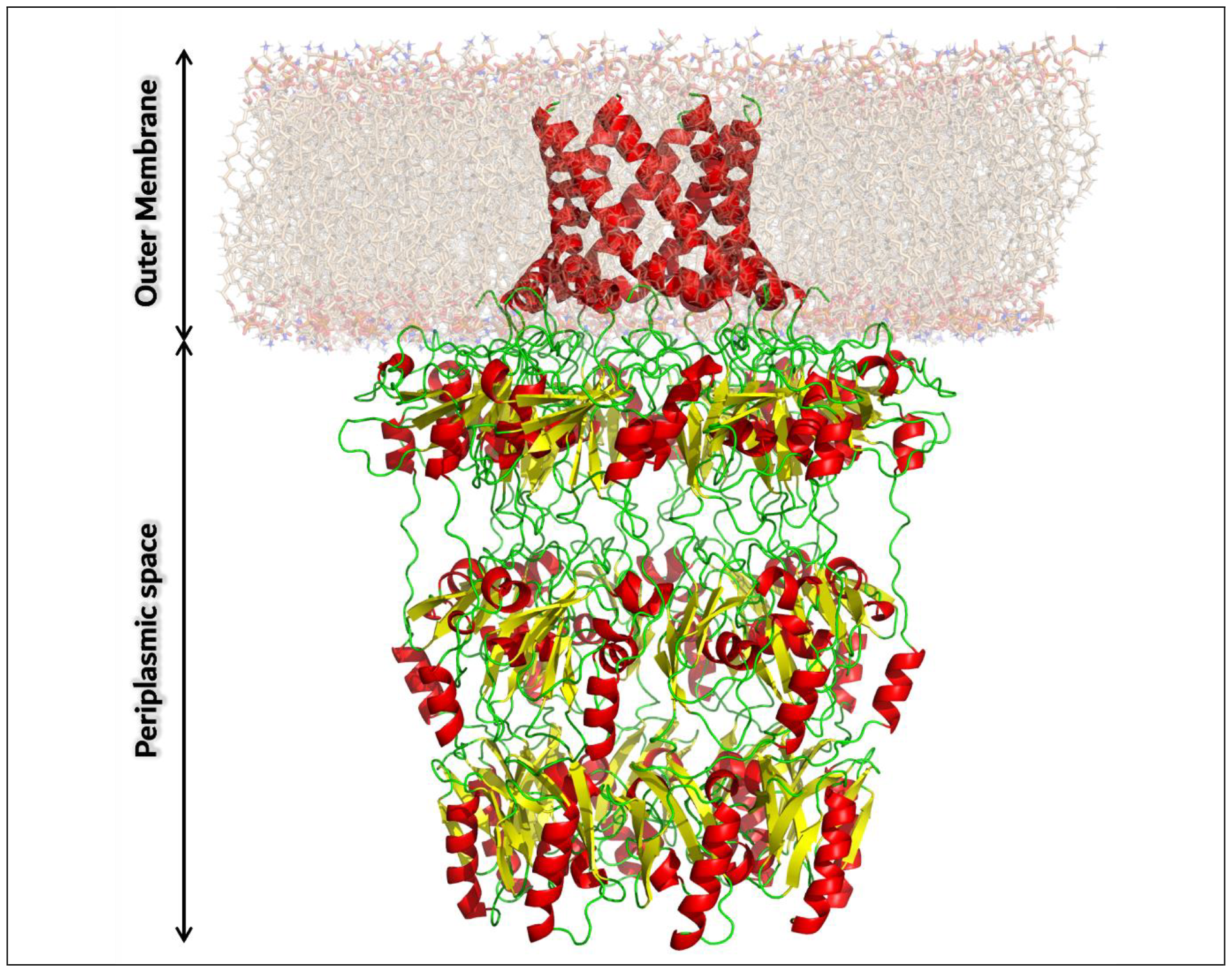
Example of a protein (*E. coli* Wza PDB ID: 2J58) that spans beyond the membrane.

**Figure S3.**
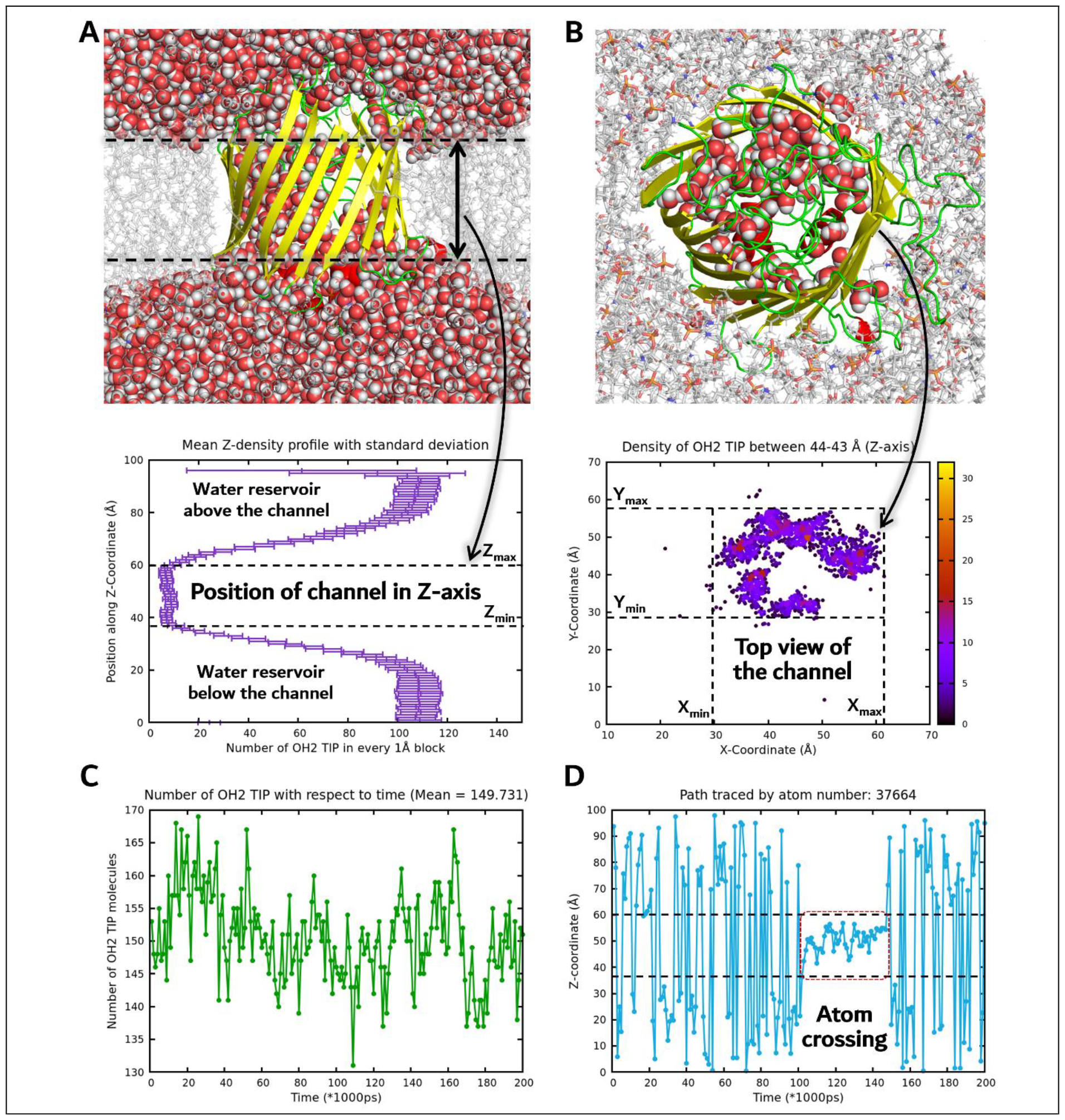
Graphical outputs generated by submodules 1, 2, 3, and 7 of APM: A) A snapshot of the MD trajectory showing the bulk water (represented by spheres) translocation across the *E*.*coli* Wzi (cartoon representation) channel: (top) front view of the channel and (bottom) the concomitant “Number of water molecules in every 1 Å block vs. Position along Z-coordinate (Å)” plot generated by the submodule “Z-density profile”. The dotted lines indicate the position of the channel in the Z-dimension. B) (top) Top view of the Wzi channel and (bottom) the corresponding position of the channel in the XY-plane projected onto the 1 Å block defined along the Z-axis (representative plot between 44 - 43 Å) generated by the “XY-area profile” module. C) The number of molecules within the channel (Z-coordinates between 40 to 60 Å as defined by Figure A (bottom)) is shown as a “Time (ps) vs. Number of molecules” plot generated by the “Rate of change of molecules” sub-module. A mean of ∼150 molecules is seen in the channel in accordance with the previous study [25]. D) The path traced by the water molecule with atom number 37664 (oxygen) is shown as a “Time (ps) vs. Z-coordinate (Å)” plot generated by the “Track molecule” module. This molecule takes about ∼50 ns (*viz*., enters and exits between ∼100-150 ns) to cross the channel from the periplasmic side to the extracellular side.

**Figure S4.**
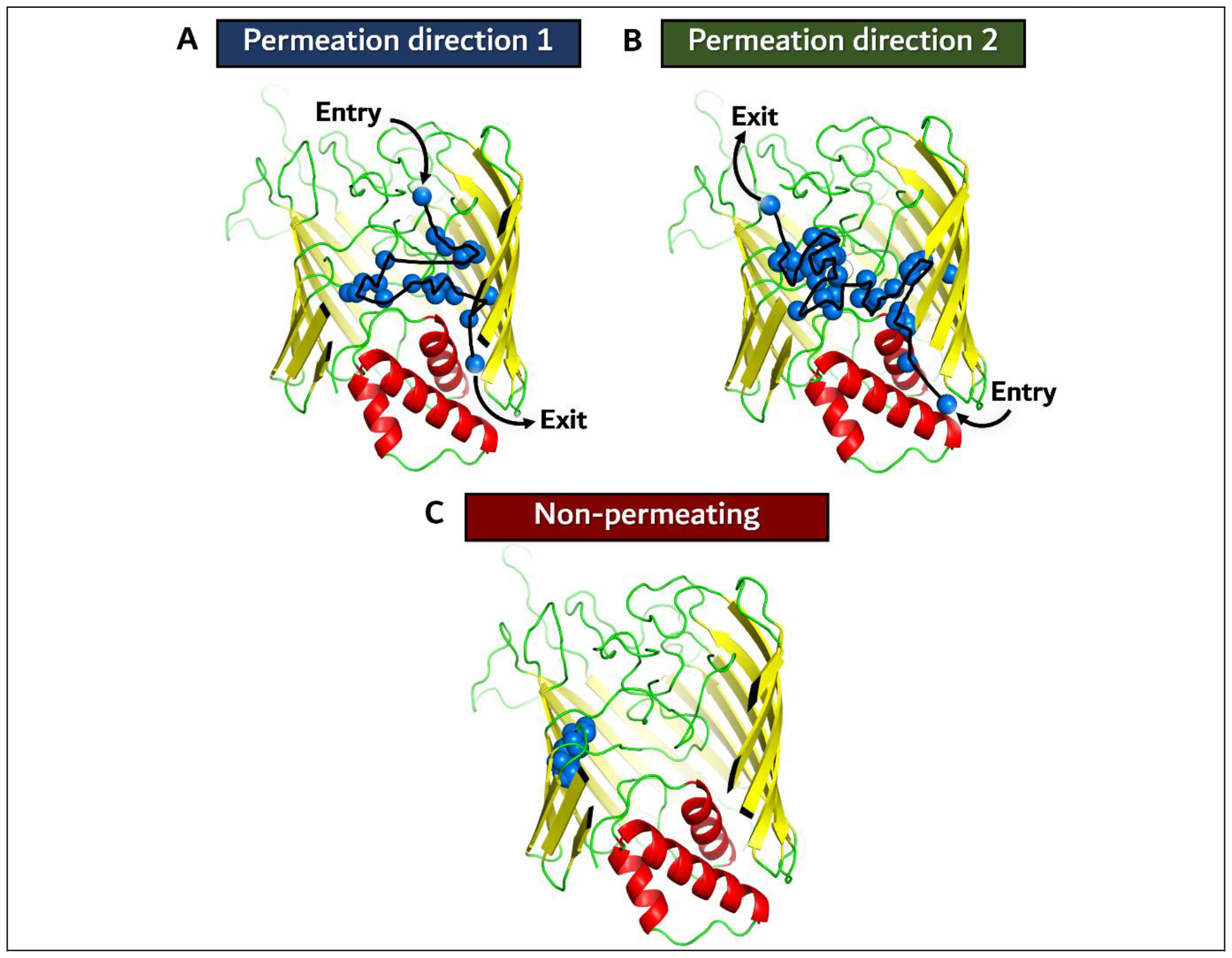
Cartoon representation of Wzi showing two permeation events: A) direction 1 and B) direction 2, along with a C) non-permeating event.

**Figure S5.**
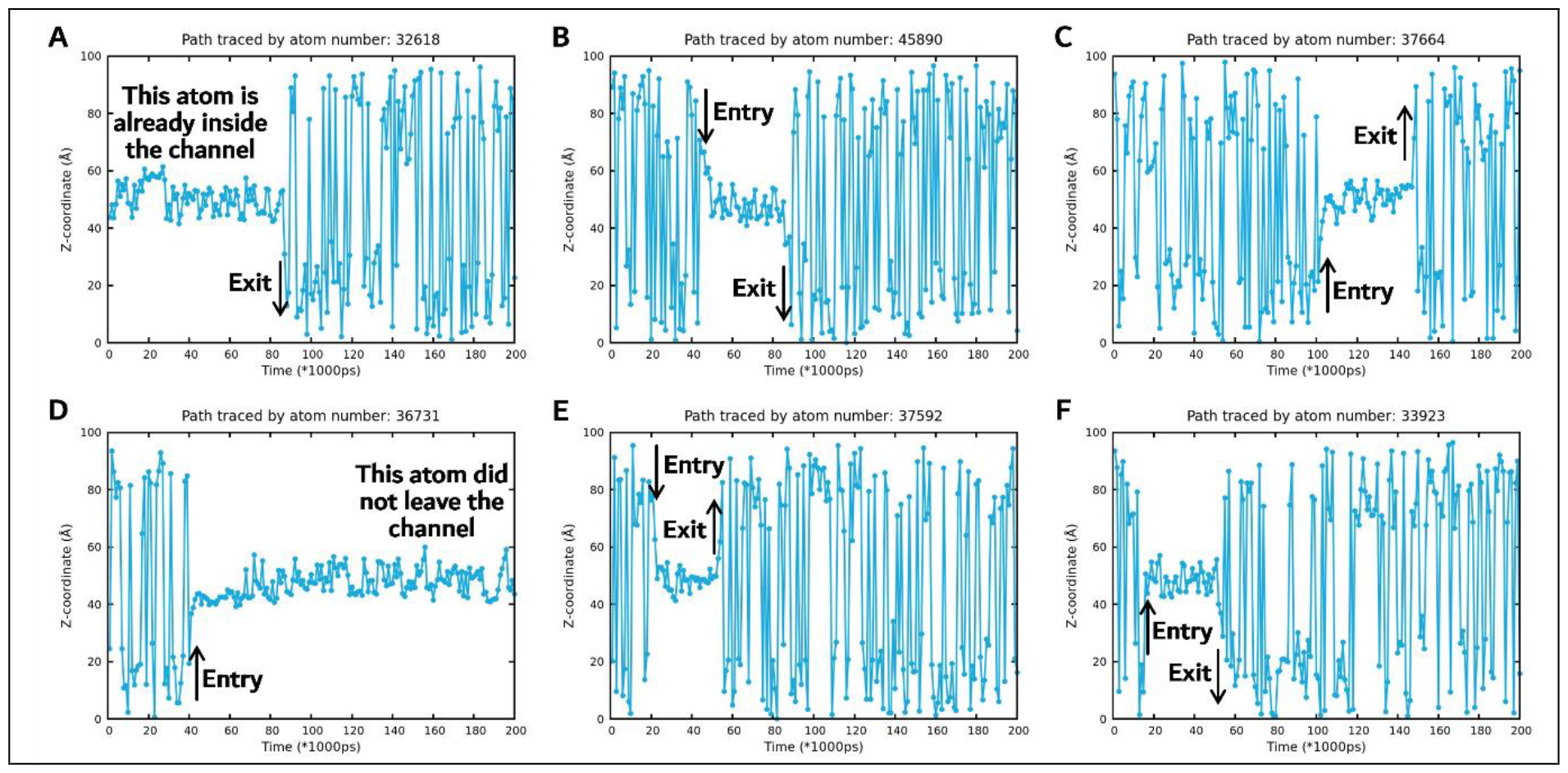
Outputs generated by the “Track molecule” module explaining the six types of permeating/non-permeating events seen during the Wzi simulation: a water molecule A) residing inside the channel at the beginning of the simulation and leaving the channel during the simulation (non-permeating), B) entering from the extracellular side and leaving from the periplasmic side (permeation direction 1), C) entering from the periplasmic side and leaving at the extracellular side (permeation direction 2), D) entering into the channel during the simulation and staying within the channel till the end of the simulation time (non-permeating), E) entering and exiting the channel from the extracellular side (non-permeating as entry and exit is from the same side), and F) entering and exiting the channel from the periplasmic side (non-permeating as entry and exit is from the same side).

**Figure S6.**
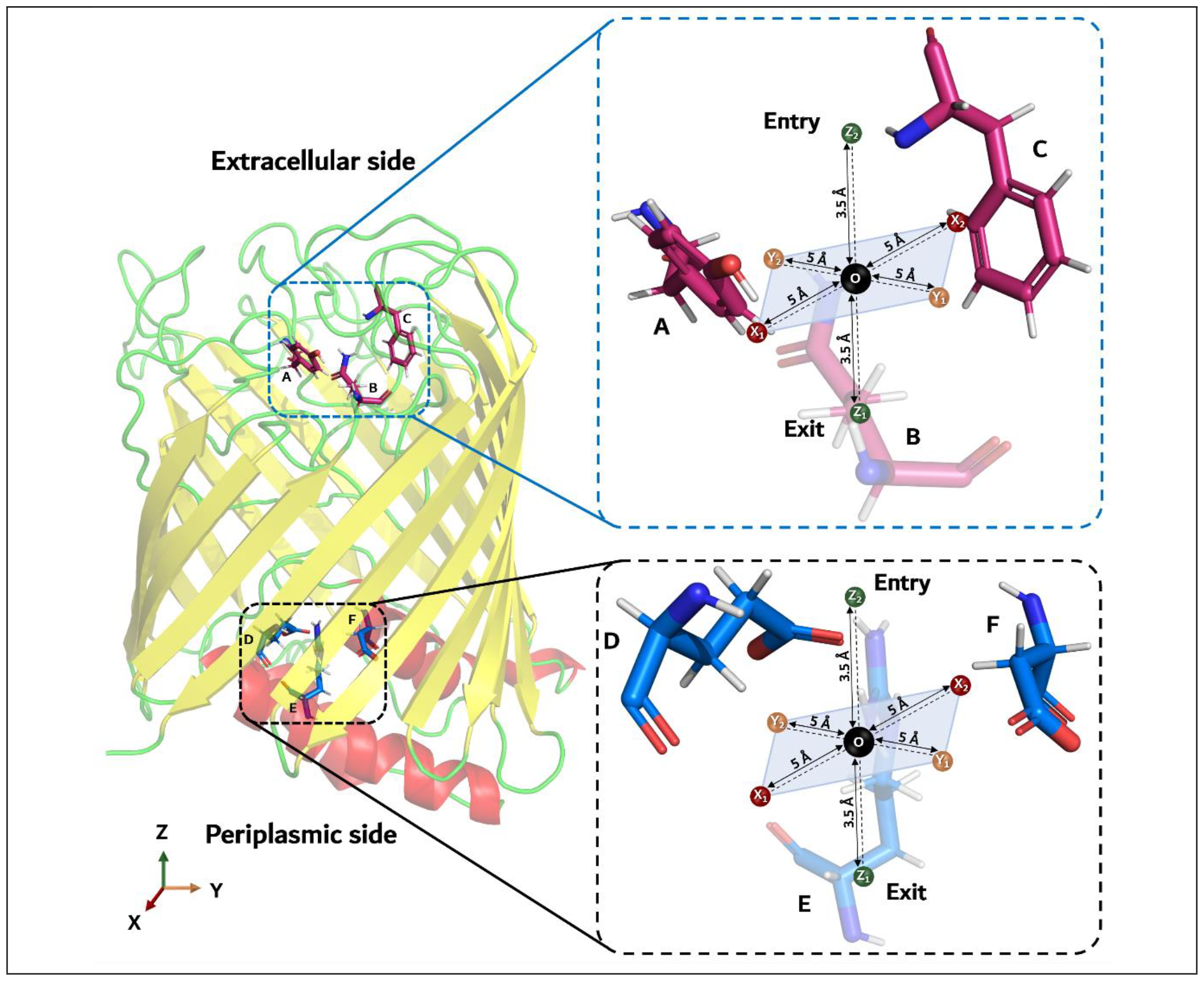
Schematic illustrating the diffusion entry and exit calculation by considering water diffusion across *E. coli* Wzi. Diffusion entry (extracellular side to the barrel) (top), exit (barrel to extracellular side) (bottom), and vice-versa are shown in the dotted box.

**Figure S7.**
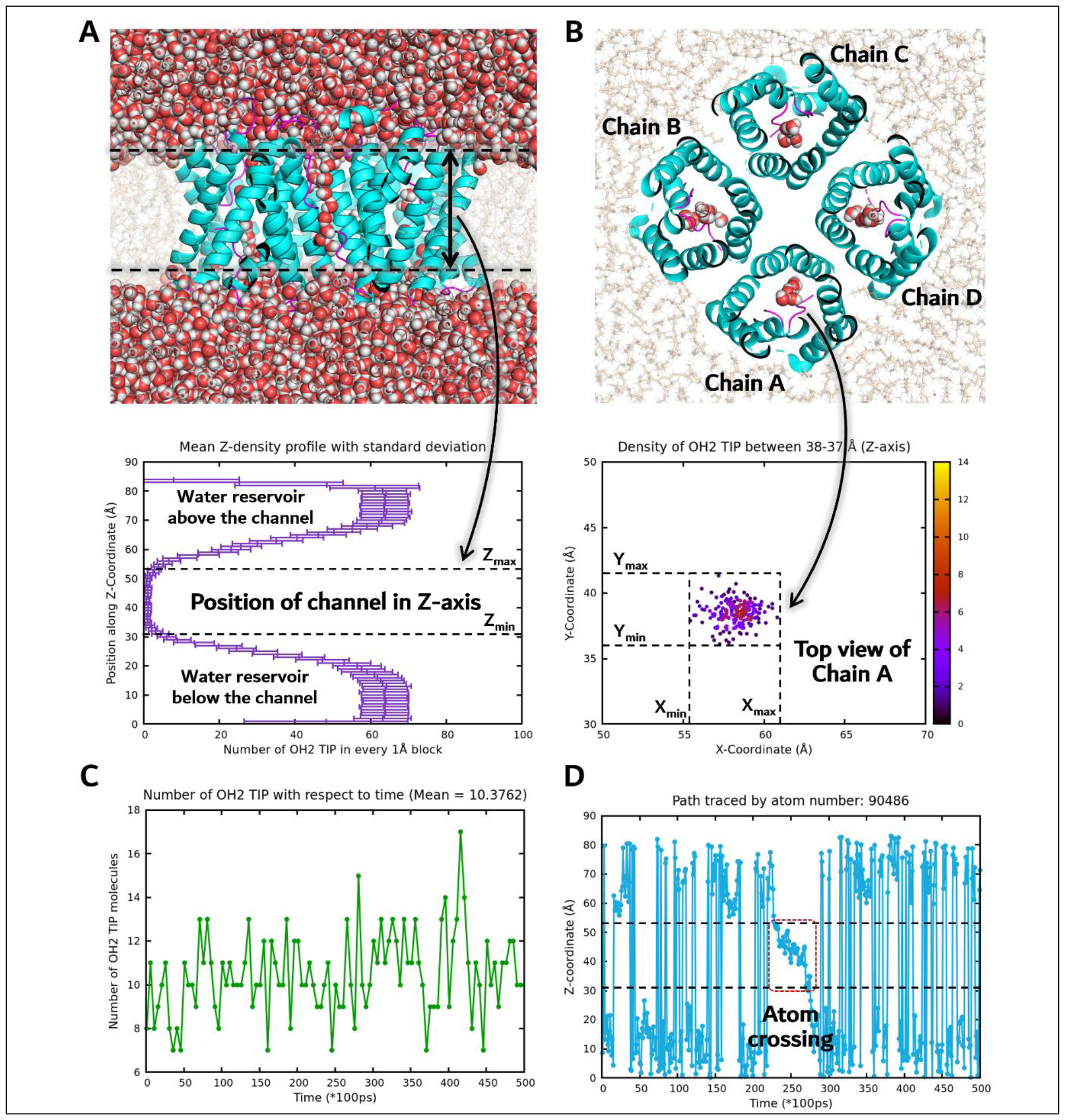
Graphical outputs generated by submodules 1, 2, 3, and 7 of APM for the 50 ns trajectory analysis of Aqp-Z (PDB ID: 2ABM): A) A snapshot of the MD trajectory showing the single file water (represented by spheres) translocation across the *E*.*coli* Aqp-Z (cartoon representation) channel: (top) front view of the channel and (bottom) the concomitant “Number of water molecules in every 1 Å block vs. Position along Z-coordinate (Å)” plot generated by the submodule “Z-density profile”. The dotted lines indicate the position of the channel in the Z-dimension. B) (top) Top view of the Aqp-Z channel and (bottom) the corresponding position of the channel in the XY-plane projected onto the 1 Å block defined along the Z-axis (representative plot between 38-37 Å) generated by the “XY-area profile” module. C) The number of molecules within the channel (Z-coordinates between 30 to 55 Å as defined by Figure A (bottom)) is shown as a “Time (ps) vs. Number of molecules” plot generated by the “Rate of change of molecules” sub-module. A mean of ∼10 molecules is seen in Chain A. D) The path traced by the water molecule with atom number 90486 (oxygen) is shown as a “Time (ps) vs. Z-coordinate (Å)” plot generated by the “Track molecule” module. This molecule takes about ∼4 ns (*viz*., enters and exits between ∼23-27 ns) to cross the channel.

**Figure S8.**
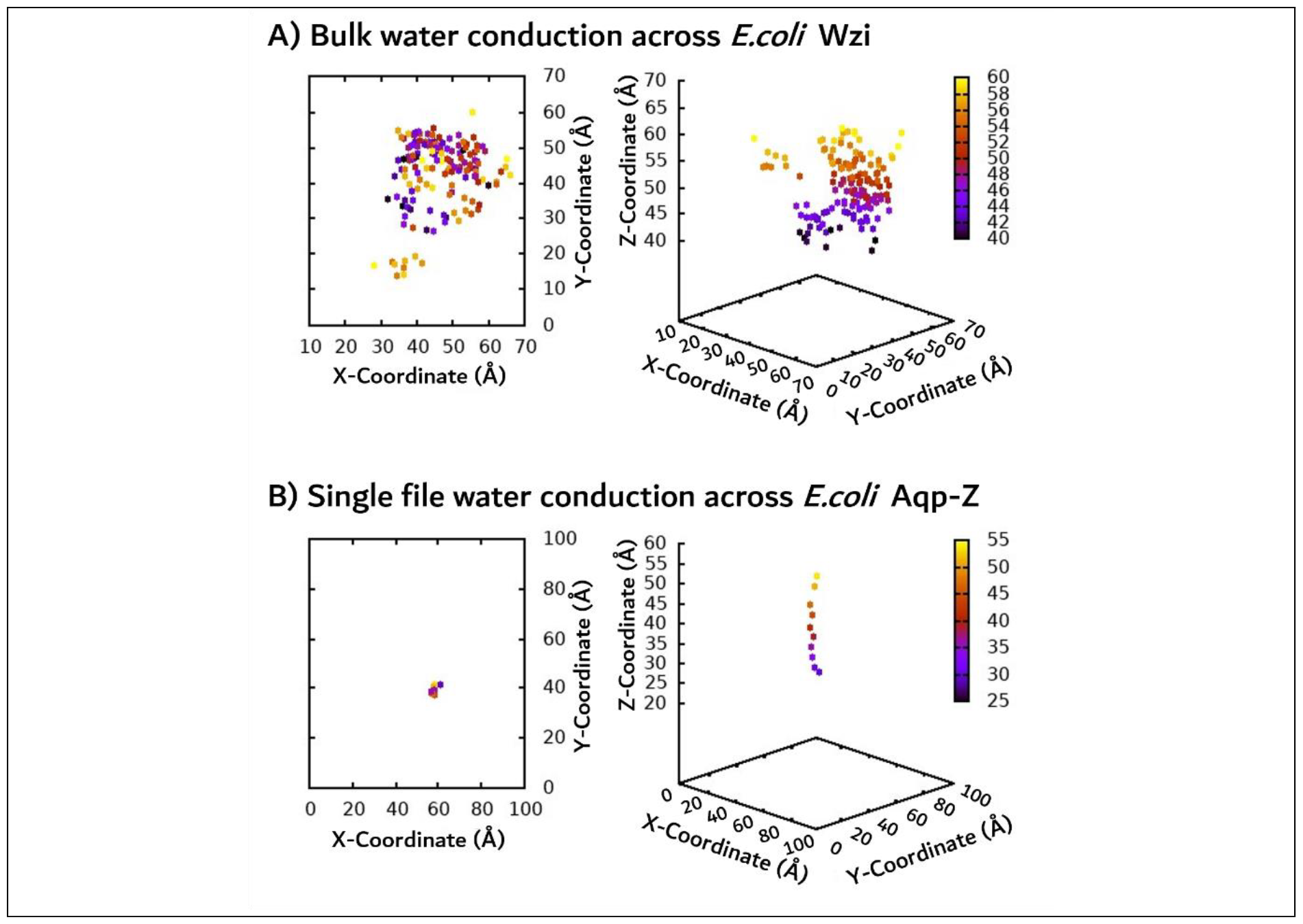
Comparison between A) bulk water conduction in *E. coli* Wzi (at 41 ns) and B) single profile conduction in *E. coli* Aqp-Z (at 6 ns): (Left) 2D and (Right) 3D plots representing the position of water molecules inside the respective channel.

**Movie 1**. Movie showing the change in the water density in the XY plane along every 1 Å Z-axis block from the top (extracellular side) of the channel to the bottom (periplasmic side): (Left) MDTAP generated plots and (Right) the corresponding representation of water conduction across Wzi protein. Note that (Left) the dots in the plots represent the density of water oxygens.

